# Exhaustion profile on classical monocytes after LPS stimulation in patients with Crohn’s disease

**DOI:** 10.1101/2024.03.28.587307

**Authors:** Lucas Pires Garcia Oliveira, Rafaela Gomes Xavier, Claudia Concer Viero Nora, Cristóvão Luis Pitangueira Mangueira, Eliane Aparecida Rosseto, Thiago Aloia, Jaime Zaladek Gil, Arceu Scanavini Neto, Filipa Blasco Tavares Pereira Lopes, Karina Inacio Carvalho

**Affiliations:** Instituto de Ensino e Pesquisa, Hospital Israelita Albert Einstein, São Paulo, Brazil; Laboratório Clínico, Hospital Israelita Albert Einstein, São Paulo, Brazil; Corpo Clínico, Hospital Israelita Albert Einstein, São Paulo, Brazil; Department of Nutrition, Case Western Reserve University, Cleveland, Ohio, USA; Case Comprehensive Cancer Center, Case Western Reserve University, Cleveland, Ohio, USA

## Abstract

Crohn’s disease is an inflammatory bowel disease that induces diarrhea, abdominal pain, weight loss, and even susceptibility to developing tumors. The immune system is pivotal in the gastrointestinal tract, promoting tolerance against commensal antigens and food. However, Crohn’s disease manifests by a breakdown in the mechanism of immune tolerance and the consequent development of exacerbated chronic inflammatory responses. The involvement of the immune system is pivotal in Crohn’s disease, with a wide range of immune cells being altered, which may include monocytes. Taking the lack of knowledge regarding monocytes in Crohn’s disease, we ought to elucidate the cytokine production and activation profile of monocyte subsets in the pathophysiology. We used multiparametric flow cytometry, quantified gene expression using qPCR, and made a correlation matrix regarding flow cytometry data and qPCR using a bioinformatic approach to examine monocyte status. The Corhn’s patients show a decrease in all subsets of monocytes.

In contrast, classical monocytes show an exhaustion profile with increased expression of CD38 and decreased production of IL-1β after LPS stimulation in the patients’ group. These results indicate that monocyte subsets are differentially involved in the pathophysiology. These findings may suggest that monocytes favor disease chronicity and lack immune response resolution.

## Introduction

Inflammatory bowel diseases (IBDs) encompass a group of chronic inflammatory conditions that affect the gastrointestinal (GI) system ^[1,2]^. Among these conditions, Crohn’s disease (CD) stands out as an important pathology that affects any part of the GI. CD is characterized by an immune response imbalance triggered by a combination of environmental, genetic, and microbiome factors. This leads to a range of symptoms in patients, including weight loss, fatigue, diarrhea, abdominal pain, and intense gut inflammation ^[3,4]^. The gut inflammation observed is mainly due to the breakdown in the immune tolerance mechanism and consequent development of inflammatory responses ^[5–8]^.

Monocytes play a vital role in the innate immune system and CD. The monocytes can be further classified into three subsets based on the expression of CD14 and CD16: classical monocytes (CD14^+^CD16^−^) with phagocytic activity and involved in innate immune response, intermediate monocytes (CD14^+^CD16^+^) responsible for cytokine secretion and antigen presentation, and non-classical monocytes (CD14^−^CD16^+^) which is associated with Fcγ-mediated phagocytosis ^[9–11]^. Classical CD14 monocytes were reported to decrease in circulating blood, suggesting their recruitment to GI-inflamed tissue ^[12,13]^. The expression of chemoreceptors CCR2 and CX3CR1 plays a crucial role in the immune response, leading to the mobilization of cells to inflamed sites and tissue homing ^[14–18]^. Once activated and recruited to the inflammation site, monocytes can release cytokines such as TNF-α and IL-1β ^[19]^. Notably, IL-1β and TNF can act as an inflammatory signal inducing lymphocyte polarization ^[20,21]^ and play an essential role in intestinal physiology ^[22]^. The other expression protein involved in anti-inflammatory signaling is CD163 ^[23]^. While CD163 is associated with tissue homeostasis and anti-inflammatory signaling, other surface markers, like CD38, correlate with cell exhaustion ^[24]^. The cellular exhaustion profile can be induced by high loads or prolonged exposure to antigens under inflammatory conditions, often observed in chronic virus infection or cancer. Cellular exhaustion eventually leads to partial or complete loss of the ability to secrete cytokines, chemokines, or degranulate. In contrast, the later exhaustion stage can induce cell death and prolong the pathology ^[25,26]^.

The study aimed to find the profile of monocytes in CD pathogenesis. Our results show a decrease of classical monocytes expressing anti-inflammatory signaling, with an increase in exhaustion profile. Future studies need to be performed to identify the functionality of the inflamed tissue.

## Material and Methods

### Cohort characterization

The Ethics Committee of the Hospital Israelita Albert Einstein approved this study (CAAE 38707914.0.0000.0071). Written informed consent was obtained from all volunteers, according to the Brazilian Ministry of Health Guidelines and Declaration of Helsinki. All samples and data were processed and stored until analysis at the Research Center of Hospital Israelita Albert Einstein.

Our cohort is composed of 210 individuals, 96 with Crohn’s patients and 114 healthy subjects. From this cohort, all 210 individuals had their buffy coat used for gene expression, using the qPCR method as subsequently described. In comparison, 67 individuals had their peripheral blood mononuclear cells (PBMC) utilized for flow cytometry assays.

### Flow cytometry

PBMCs from 26 Crohn’s patients and 41 healthy subjects were thawed and washed in FACS buffer (PBS 1x, FBS 1%); viability was assessed using a Countess automated cell counter (ThermoFisher Scientific, USA). Staining was performed for 30 min at room temperature in 96-well V bottom plates (Thermo-Fisher Scientific, USA) with the following antibodies against CD3 (clone: SK7), CD11b (clone: ICR44), CD14 (clone: M5E2), CD15 (clone: W6D3), CD16 (clone: 3G8), CD33 (clone: P67.6), CD38 (clone: HB-7), CD163 (clone: GHI/61), CCR2 (clone: K036C2), CX3CR1 (clone: 2A9-1) from BioLegend. Amine Aqua dye (ThermoFisher Scientific, USA) was used to exclude dead cells. To measure the best cytokine production, we performed kinetics to detect IL-1β and TNF-α in frozen PBMCs. PBMCs were thawed in the presence of 100 ng/ml LPS (Sigma-Aldrich, USA) and incubated at 37°C and 5% CO_2_ for 1, 2, 4, 6, and 8 hours. In 30 minutes of incubation, monensin (5 mg/ml; Golgi Stop, BD Biosciences, USA) and Brefeldin A (5 mg/ml; Golgi Plug, BD Biosciences, USA) were added to the medium for the 1-hour point. After 2, 4, 6, and 8 hours of incubation, monensin and Brefeldin A were added after 1 hour of incubation. After incubation for 2 h, cells were washed, and surface staining was performed at room temperature for 15 min. Cells were fixed/permeabilized using the Fix & Perm Cell Permeabilization kit from Life Technologies. Cells were incubated with antibodies against TNF-α (clone: Mab11) and IL-1β (clone: CRM56) from Biolegend for 60 minutes. Samples were washed and fixed with formaldehyde before flow cytometry data acquisition.

The best secretion point of both cytokines was at 2 hours incubation (S1 Fig), and all samples were performed at this time point. One pitfall of such a technique for monocytes is the expression of CD16; the monocytes lose their surface expression after 2 h of incubation with LPS, as already demonstrated ^[27]^; thus, we were not able to assess the cytokine production by the intermediate and non-classical monocyte subsets.

The Fluorescence minus one (FMO) ^[28]^ was used as a gating strategy for surface panels, and unstimulated cells were used for intracellular analysis. All samples were acquired using an LSR Fortessa flow cytometer (BD Biosciences, USA) and FACSDiva software (BD Biosciences, USA). Gate analysis was performed using the FlowJo software version 10.9 (BD Biosciences, USA).

### RNA extraction

RNA was extracted from 96 Crohn’s patients and 114 healthy buffy coat samples using the Purelink RNA Minikit, described by the manufacturer (Life Sciences, USA). After digestion, residual DNA was removed using DNase A. RNA was eluted from the column using a low-salt solution buffer. NanoDropC (Invitrogen, USA) measured the concentration and RNA integrity.

### Real-time quantitative PCR (RT-qPCR)

Quantitative polymerase chain reaction (qPCR) was performed using a QuantStudio^®^ 6 real-time PCR system to assess changes in mRNA expression in the genes (S1 Table). Total RNA was extracted from cells using a PureLink RNA mini kit (Life Sciences, USA). cDNA synthesis was performed with a High-Capacity cDNA Reverse Transcription Kit (Life Sciences, USA) according to the manufacturer’s instructions. qPCR was performed on a total of 20 μl containing Quantinova SYBR™ Green PCR kit (Quiagen, DE) according to manufacturer instructions. The relative expressions were analyzed by the comparative CT method (ΔΔCt), normalized by the expression of β-actin and GAPDH, comparing the static conditions as a reference.

**Supplementary Table 1.**
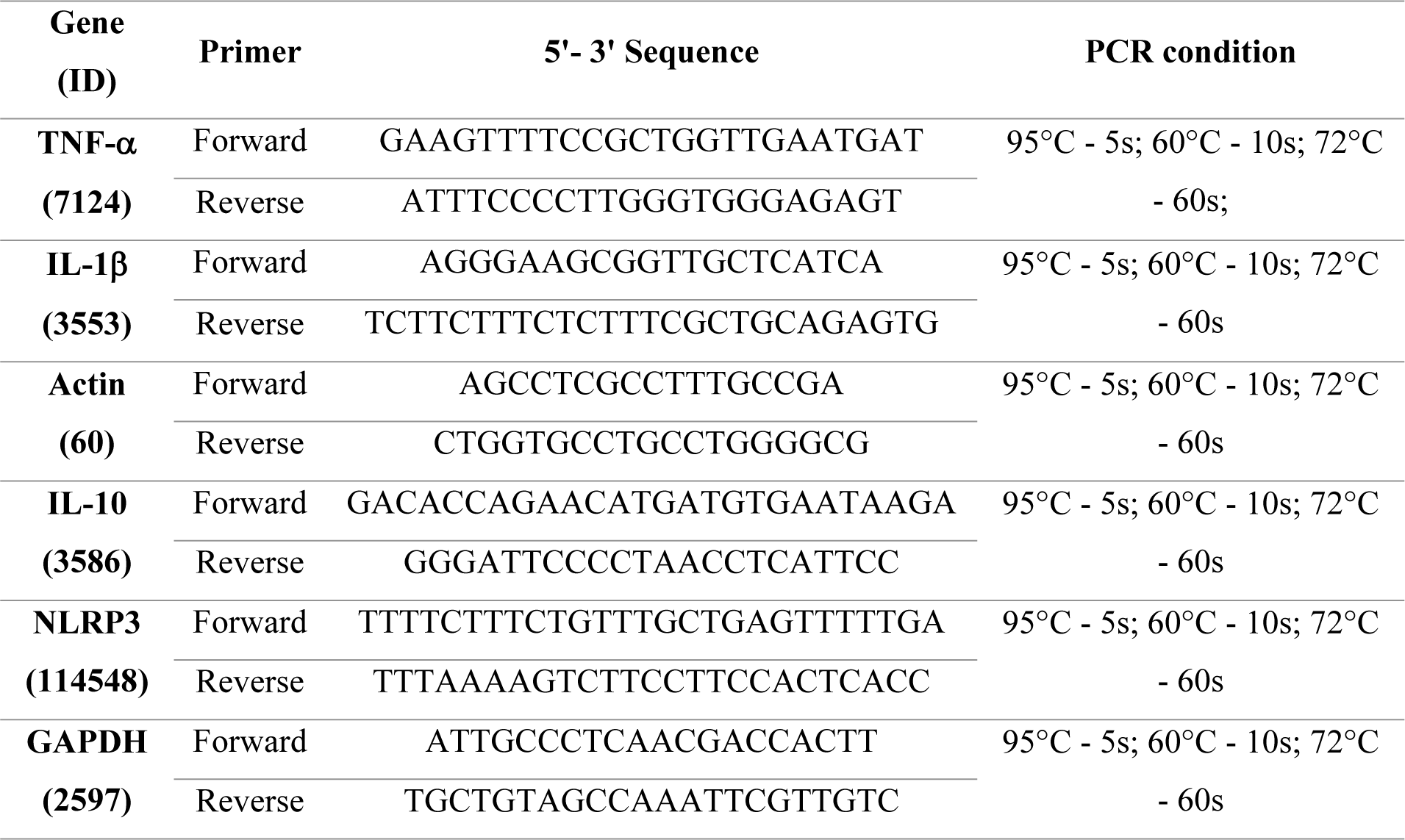
Primer sequence and PCR conditions.

### Immunofluorescent Staining

Colon sections were cut 5mM, after heat-induced epitope retrieval, cells were incubated overnight at 4°C with rat anti-human HLA-DR (clone: YD1/63.4) monoclonal antibody (Thermo-Fisher Scientific, USA), rabbit anti-human CD16 (clone: JE49-79) polyclonal antibody (Invitrogen, USA) and mouse anti-human CD64 (clone: OTI3D3) monoclonal antibody (Abcan, UK). After washing, sections were stained with a secondary antibody anti-rat Alexa 647 (Thermo-Fisher Scientific, USA), anti-rabbit Alexa 568 (Thermo-Fisher Scientific, USA), and anti-mouse Alexa 488 (Thermo-Fisher Scientific, USA) for two hours at room temperature. The sections were washed and incubated for two hours with anti-human mouse CD14 Texas Red (clone: RMO52) (Beckman Coulter, USA). Afterward, the sections were incubated with DAPI and mounted on slides using Prolong Gold antifade reagent (Thermo-Fisher Scientific, USA). Images were acquired on an LSM 710 confocal microscope (Carl Zeiss, DE) with a 20X objective. The colocalization was assessed by using Zen 2012 SP2. The expression of HLA-DR^+^CD64^+^ cells defined the monocyte-like cells compartment. Ten different fields were counted for the 5 lesion and 6 margin sites.

### Statistical analysis

Statistical analysis was performed using Prism8 (GraphPad Software, USA), considering a significance level of 5% (p ≤ 0.05), and represented in the figures are the median, 1^st^, and 3^rd^ interquartile values (M: Q1-Q3). Mann-Whitney was used to compare the cohort characteristics between the groups, and the Given Proportions test was used to assess the differences in women’s and men’s proportions (Table 1). For the correlation matrix, were used the R package Corrplot (v. 0.92), and Pearson’s correlation was used to access each possible correlation between flow cytometry and qPCR data, considering a significance level of 5% (p≤0.05); other arguments were left default.

### Results

#### Study volunteers

The cohort is primarily women, with no differences in ratios (p>0.05), with a mean age of 33 years for the healthy group and 36 years for the Crohn’s group. It is noteworthy that the serological markers commonly used for CD, calprotectin, ASCA IgG, and ASCA IgA were significantly increased (p<0.0001) in the Crohn’s group when compared to healthy individuals (Tab 1). Furthermore, the total count of cells did not show significant differences in total leukocytes and monocyte numbers. The mean age of patients in the present study reflects a relatively young population affected by CD, as global data show that the most affected age group is above 50 ^[29]^. The average age of patients observed in Brazil may be related to the Brazilian age profile, which is mainly composed of people under 34 years old ^[30]^.

**Table 1:**
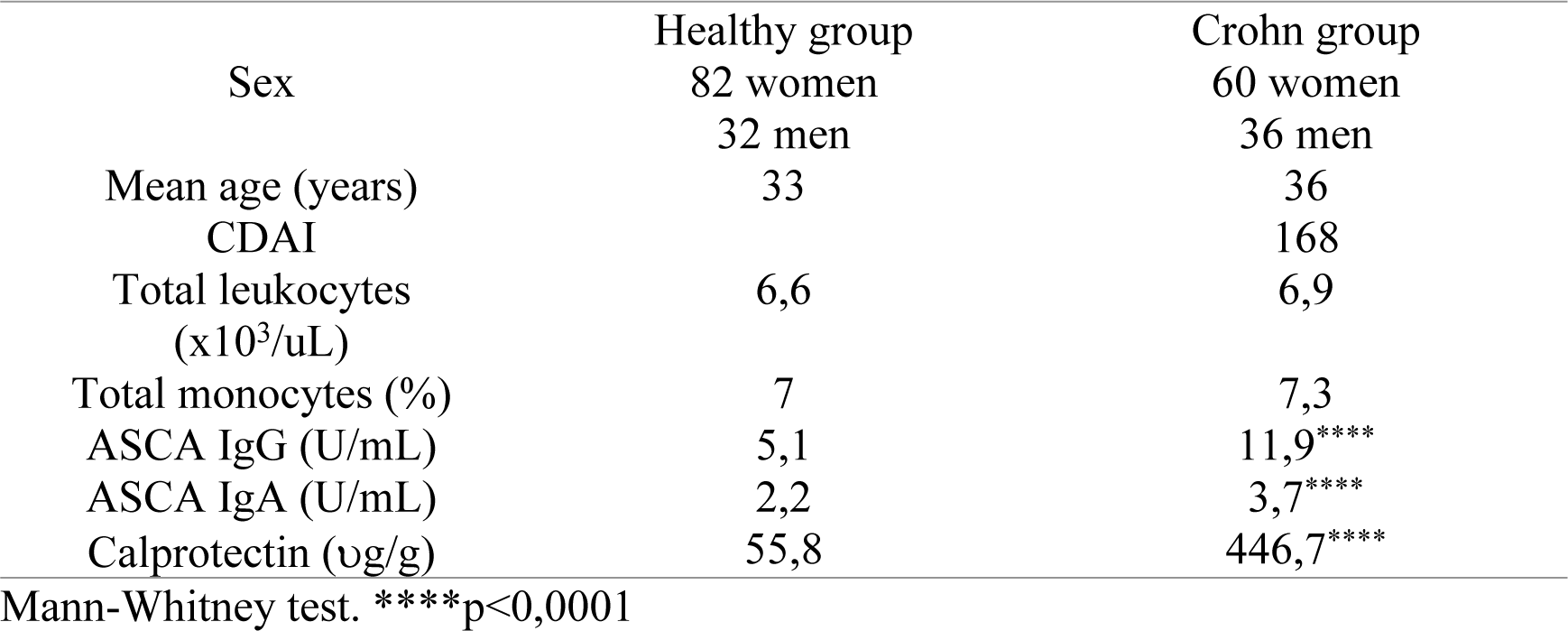
Cohort characterization.

| Sex | Healthy group<br>82 women<br>32 men | Crohn group<br>60 women<br>36 men |
| --- | --- | --- |
| Mean age (years) | 33 | 36 |
| CDAI |  | 168 |
| Total leukocytes<br>(x10<sup>3</sup>/uL) | 6,6 | 6,9 |
| Total monocytes (%) | 7 | 7,3 |
| ASCA IgG (U/mL) | 5,1 | 11,9**** |
| ASCA IgA (U/mL) | 2,2 | 3,7**** |
| Calprotectin (ug/g) | 55,8 | 446,7**** |

Mann-Whitney test. \*\*\*\*p<0,0001

### Classical Monocytes are decreased in Crohn’s patients

After the cohort was characterized, we performed the multiparametric flow cytometry assay of samples from patients (n = 26) and healthy subjects (n = 41) to measure the percentage of each monocyte subset. A representative gate strategy is represented in Fig 1A, and gate strategy of monocyte subsets on S2 Fig.

**Figure 1.**
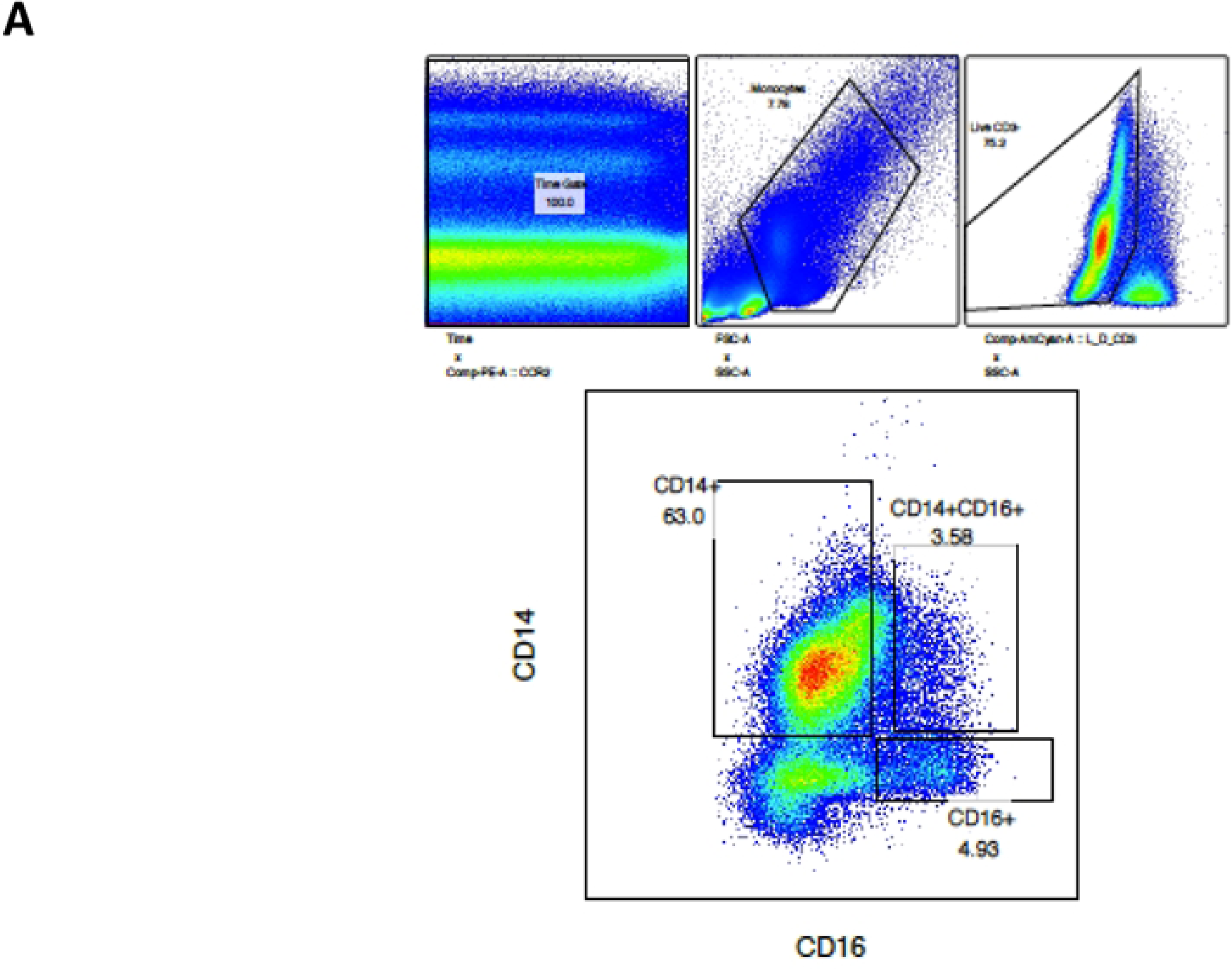
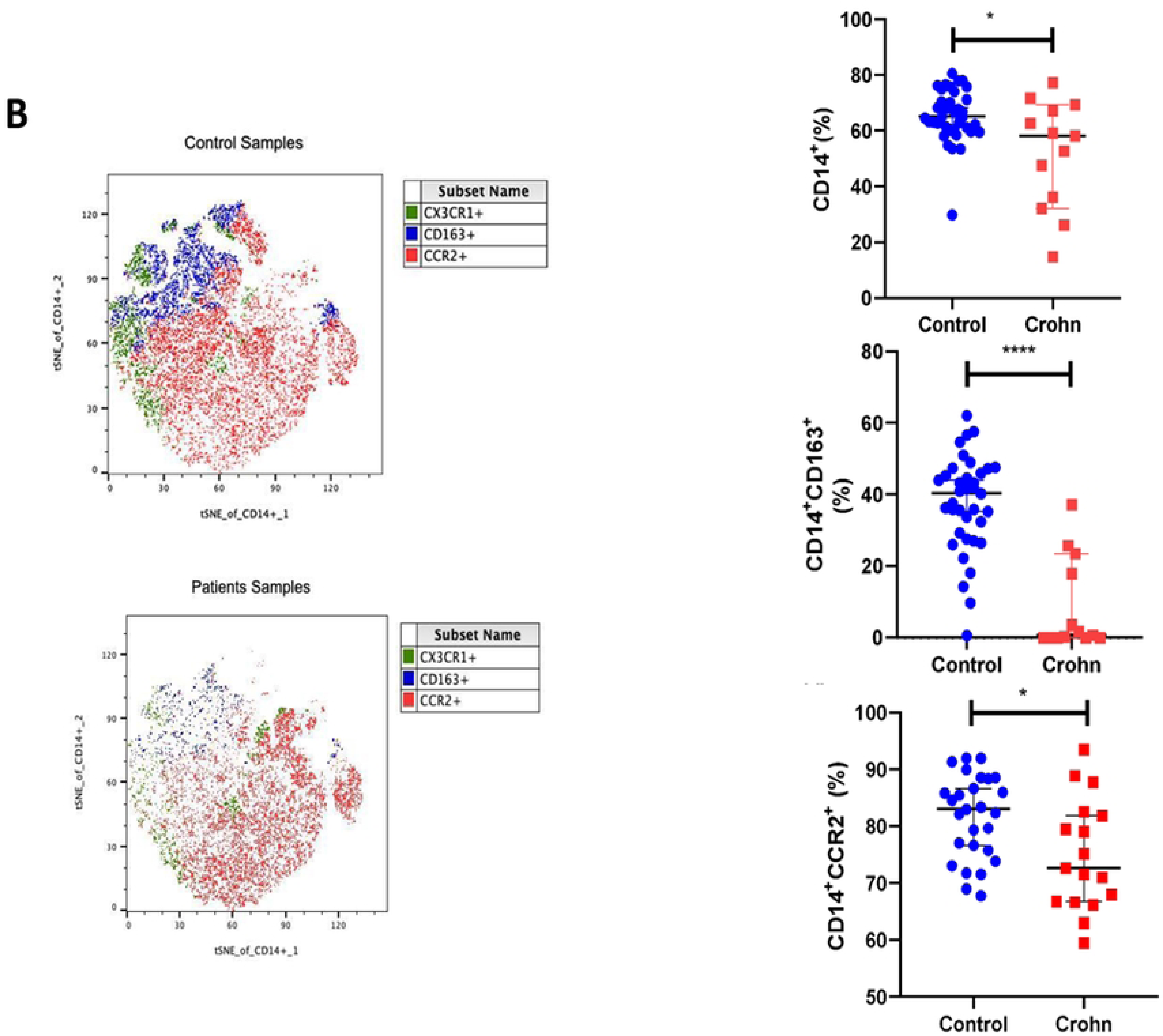
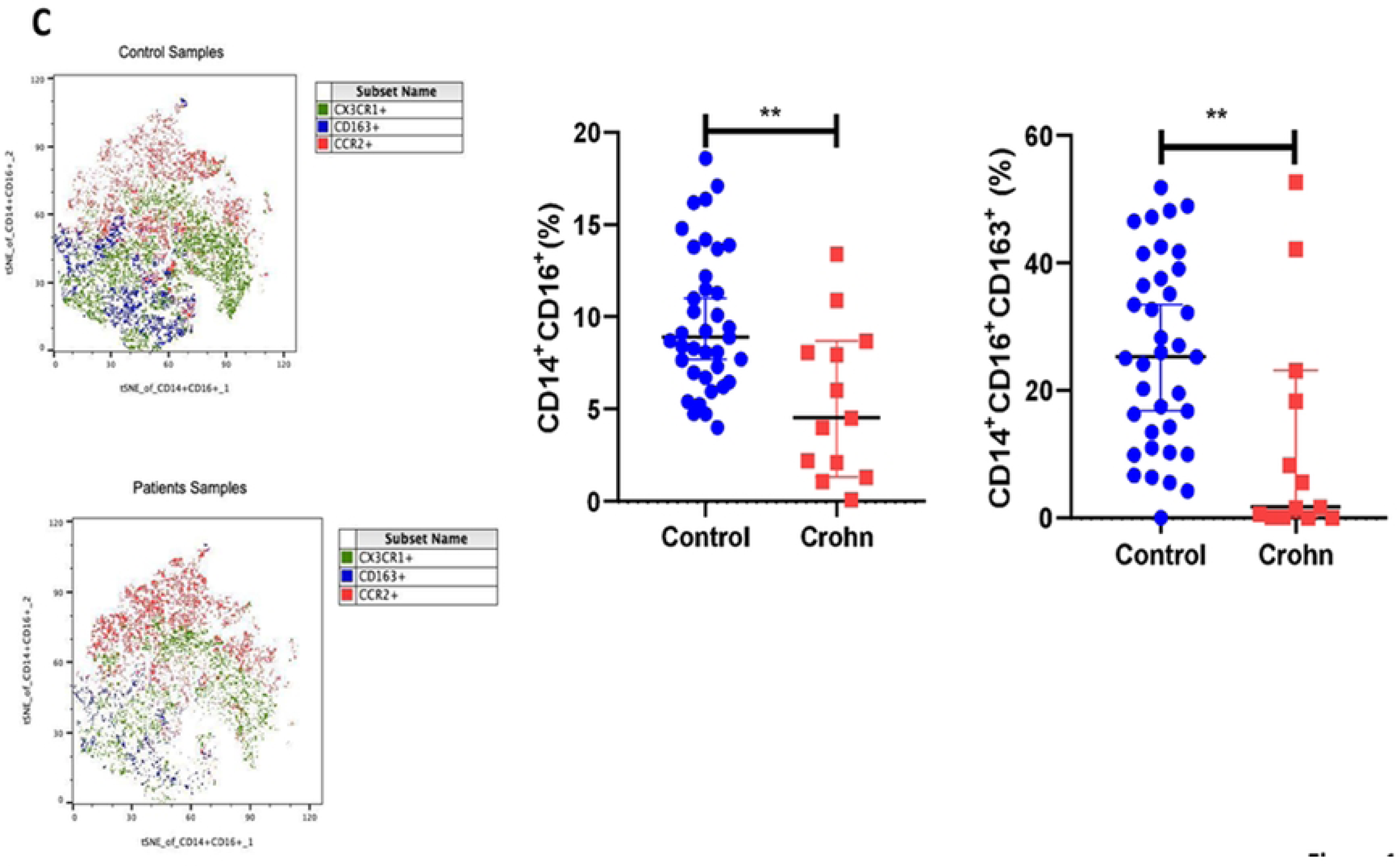
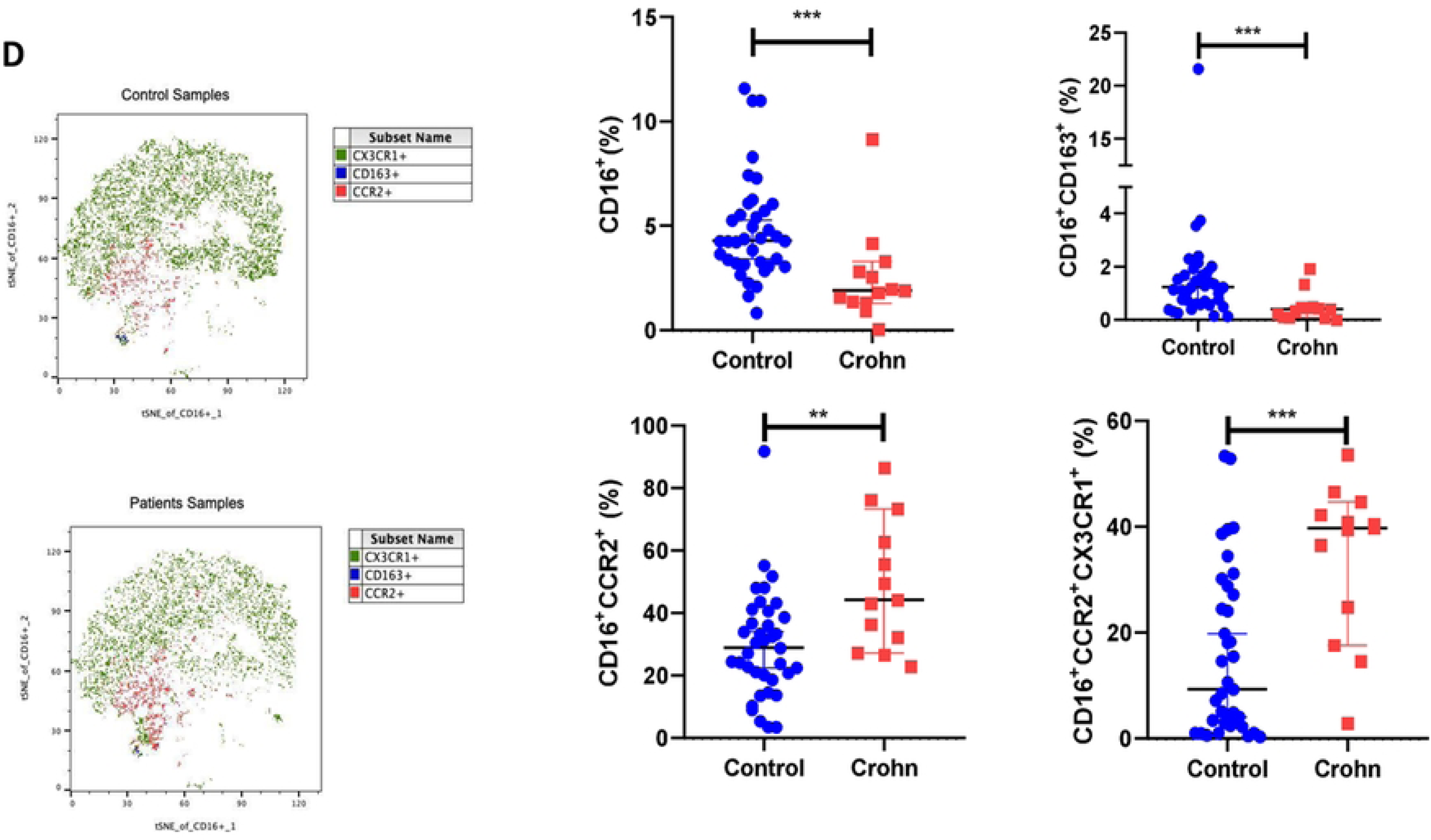
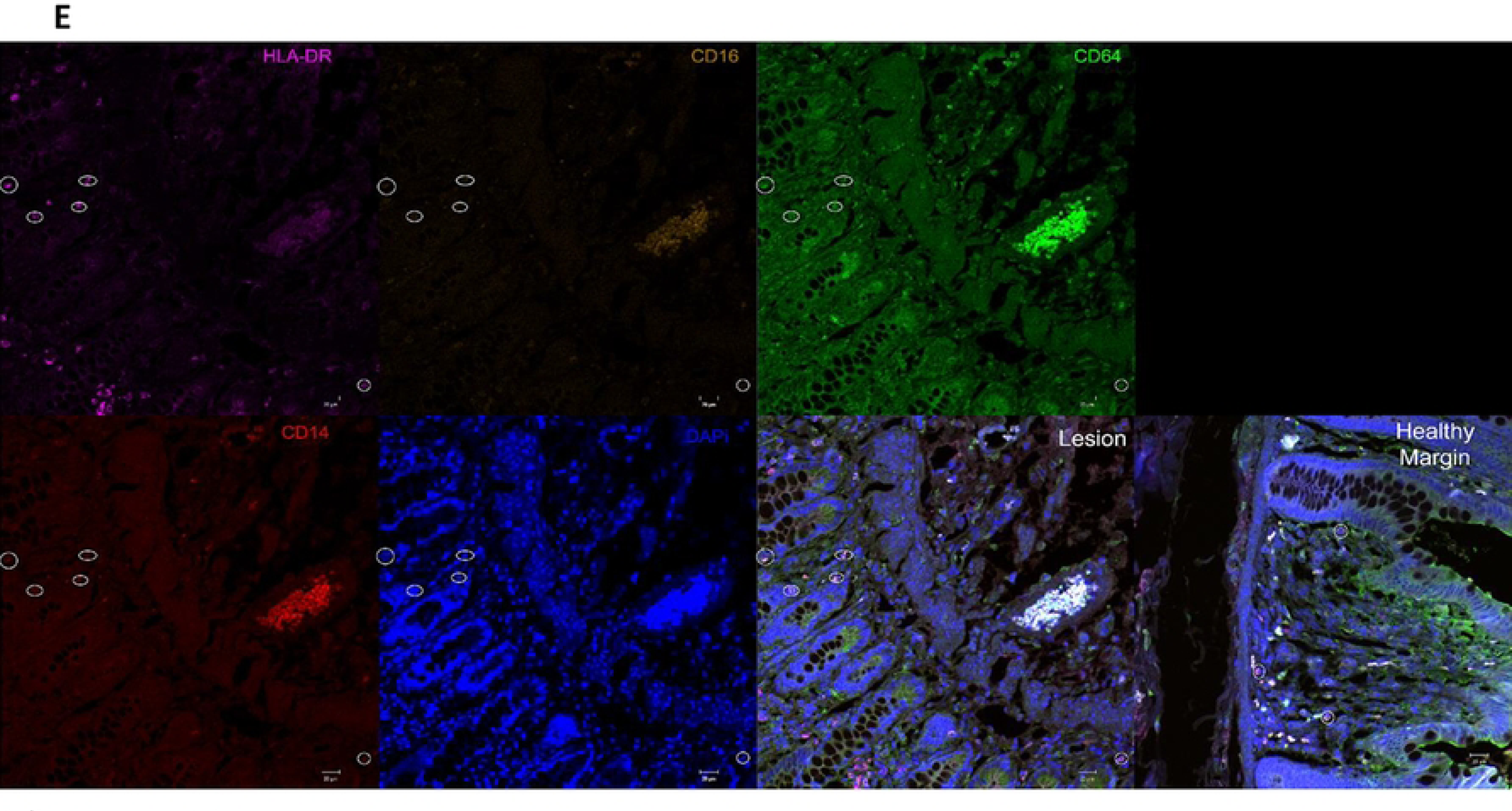
Surface markers profiles of monocytes in Crohn’s patients and healthy subjects. **(A)** Flow Cytometry gating strategy to identify the monocyte subsets. **(B)** Phenograph-based visualization on the tSNE plot of (from left to right) classical monocytes and expression of CD163 and CCR2. **(C)** tSNE plot of (from left to right) intermediary monocytes and expression of CD163. **(D)** tSNE plot of (from left to right) non-classical monocytes and CD163, CCR2, and CCR2/CX3CR1 expression. Annotation of each subcluster on merged tSNE CX3CR1 are green, CD163 blue, and CCR2 red plots, respectively. **(E)** Confocal microscopy shows a lesion and a healthy margin.

Cluster assignments show clearly that the healthy subjects show more expression of the subset markers on classical monocytes when compared with Crohn’s patients, showing in the graphics for the classical monocytes (CD14^+^CD16^−^) (p<0.05), classical monocytes expressing CD163 (CD14^+^CD16^−^CD163^+^) (p≤0.001), and classical monocytes expressing CCR2 (CD14^+^CD16^−^ CCR2^+^) (p≤0.05). Our results demonstrate a decrease in monocytes in the blood, which shows a chronic inflammatory profile and a homing of these cells for the tissue lesions represented by CCR2 (Fig 1B).

### Decreased anti-inflammatory receptor on intermediate monocytes in Crohn’s patients

CD163 protein is a surface receptor with anti-inflammatory action, essential for intestinal homeostasis. We observed diverse expressions on cluster assignments between healthy controls and Crohn’s patients. However, the statistical significance was detected in an increased percentage of intermediated monocytes (CD14^+^CD16^+^) (p≤0.01) and intermediated monocytes expressing CD163 (CD14^+^CD16^+^CD163^+^) (p≤0.01) on healthy subjects when compared with Crohn’s patients (Fig 1C). This decrease in the expression of intermediate monocytes suggests an imbalance in the activation and proliferation of T cells since this subset of monocytes is one of the immune cells responsible for this function ^[31]^.

### Chemokine receptor expression increased in non-classical monocytes

Non-classical monocytes (CD14^−^CD16^+^) and these cells expressing CD163 (CD14^−^CD16^+^CD163^+^) show a decrease in Crohn’s patients when compared with healthy subjects (p≤0.01). Then, we evaluated the expressions of CCR2 and CX3CR1 on these cells. The cluster assignments show that Crohn’s patients have CCR2 with more intensity on the cluster. The non-classical monocytes show an increase of CCR2 (CD14^−^ CD16^+^CCR2^+^) (p≤0.01) and the double expression of CCR2 CX3CR1 (CD14^−^ CD16^+^CCR2^+^CX3CR1^+^) (p≤0.001) in Crohn’s patients when compared with healthy subjects. These results show these cells’ capacity to transmigrate to injured tissue ^[32]^ (Fig 1D).

### No monocyte migration to the tissue of Crohn’s patients

Since we observed an increase in chemokine receptors, we decided to investigate the possibility of infiltration of monocytes in the injured tissue. Our imaging analysis showed no difference in the presence of cells expressing HLA-DR^+^CD64^+^ when comparing lesion tissue and disease-free margins. Unfortunately, we had only six samples of patients, and this could impact the results. Another fact could be that other types of leucocytes, such as neutrophils, accumulate in the acute lesions (Fig 1E) ^[33]^. However, active human colon IBD was associated with mRNA expression of HLA-DR and CD14^[34]^.

### Patients with Crohn’s disease show higher numbers of immature myeloid cells

In addition to monocytes, Myeloid-Derived Suppressor Cells (MDSC) may play a role in the pathogenesis of CD. To better describe these cells, we use the following makers (CD14^−^CD33^+^CD15^+^CD11b^+^) to define MDSC, as demonstrated by the gating strategy in Fig 2A. Our results showed no significant difference (*p*>0.05) between the health and patient groups despite MDSCs having an important role in chronic inflammatory pathologies and not seeming to take part in our cohort of CD in this work.

**Figure 2.**
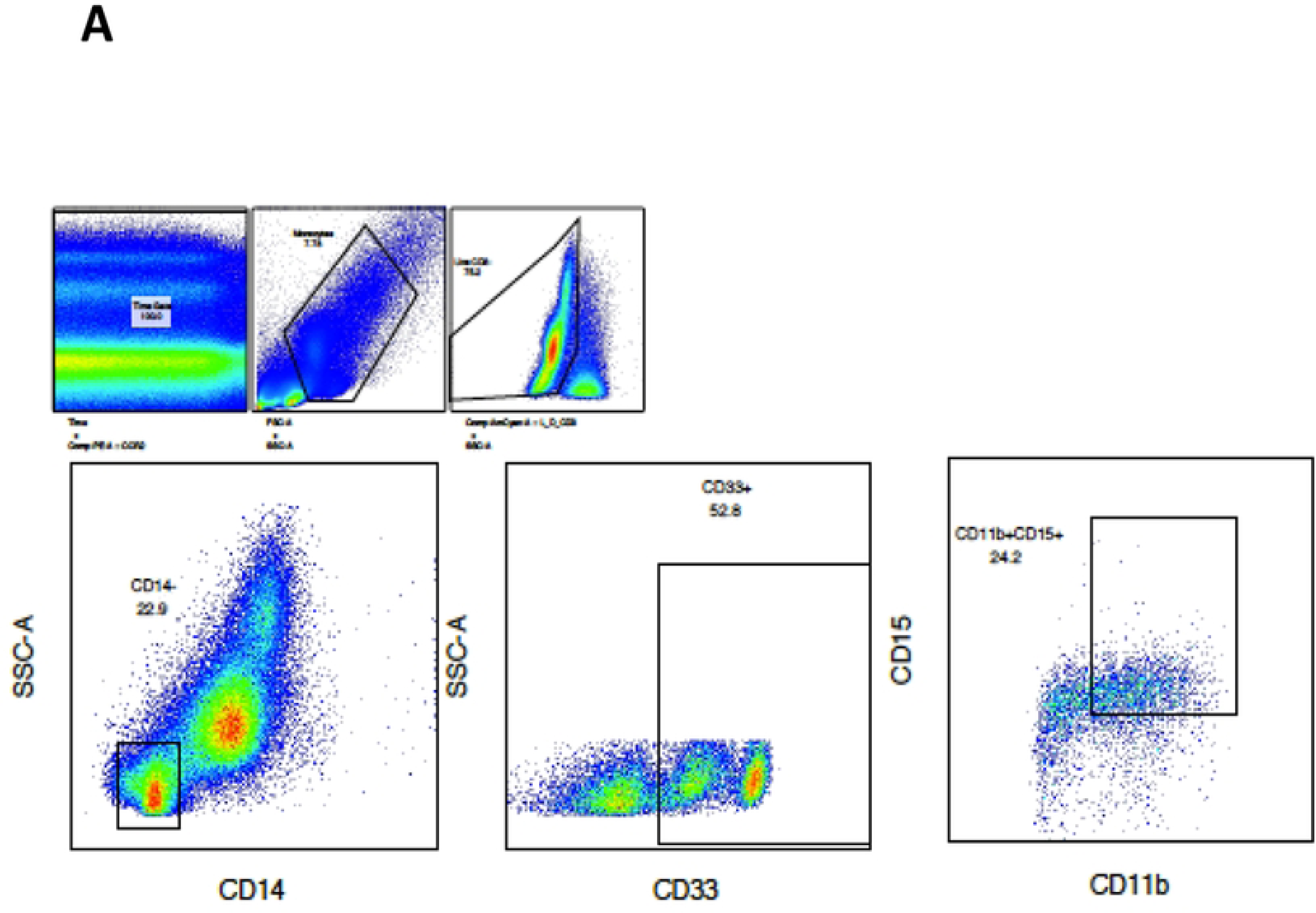
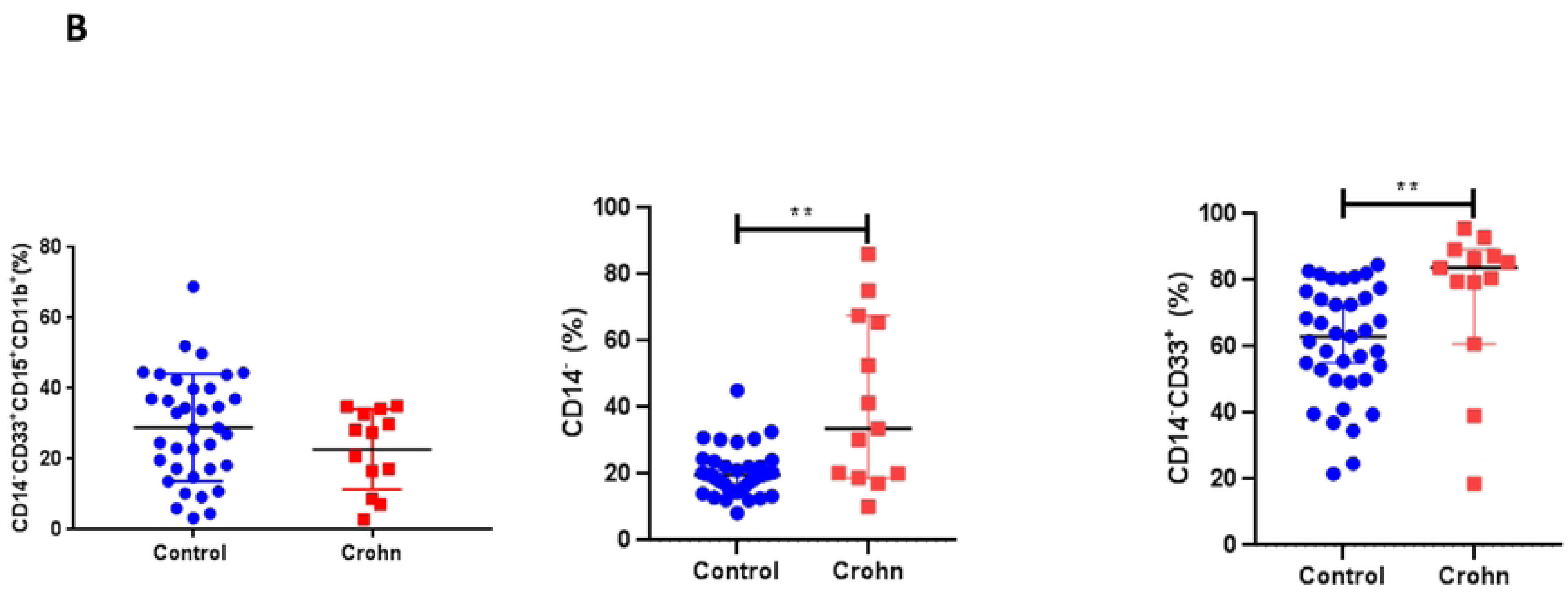
Myeloid-Derived Suppressor Cells and Myeloid immature cells in Crohn’s disease. **(A)** Representative gating strategy to identify the myeloid-derived suppressor cells and myeloid immature cells. **(B)** From left to right, the percentage of MDSC cells showed no significance, with CD14 negative cells and immature myeloid cells showing an increase in Crohn’s.

However, we observed an increase in CD14^−^ and CD14^−^CD33^+^ in Crohn’s patients when compared with health subjects (p≤0.01) (Fig 2B). The difference in CD33^+^ cells may indicate an increase in immature cells in the peripheral blood of Crohn’s patients, which may play an important role in the disease resolution not being able to build a proper immune response ^[35,36]^.

### LPS stimulus decreases chemokine receptor expression while increasing CD38 expression in classical monocytes

After incubating the PBMC of CD patients and health subjects for 2 h with LPS, as described in the methods section, we observed differences in the cytokine production and expression of surface molecules.

The percentage of CD14^+^CCR2^+^ in Crohn’s patients was decreased (p<0.05) compared to the healthy subjects. On the other hand, CD14^+^CCR2^+^CD163^+^, CD14^+^CX3CR1^+^ CD38^+,^ and CD14^+^CX3CR1^+^CCR2^+^ were increased (*p*<0.05) in CD patients in comparison to healthy subjects (Fig 3A). These results show a profile of exhaustion and the inflammatory process of classical monocytes.

**Figure 3.**
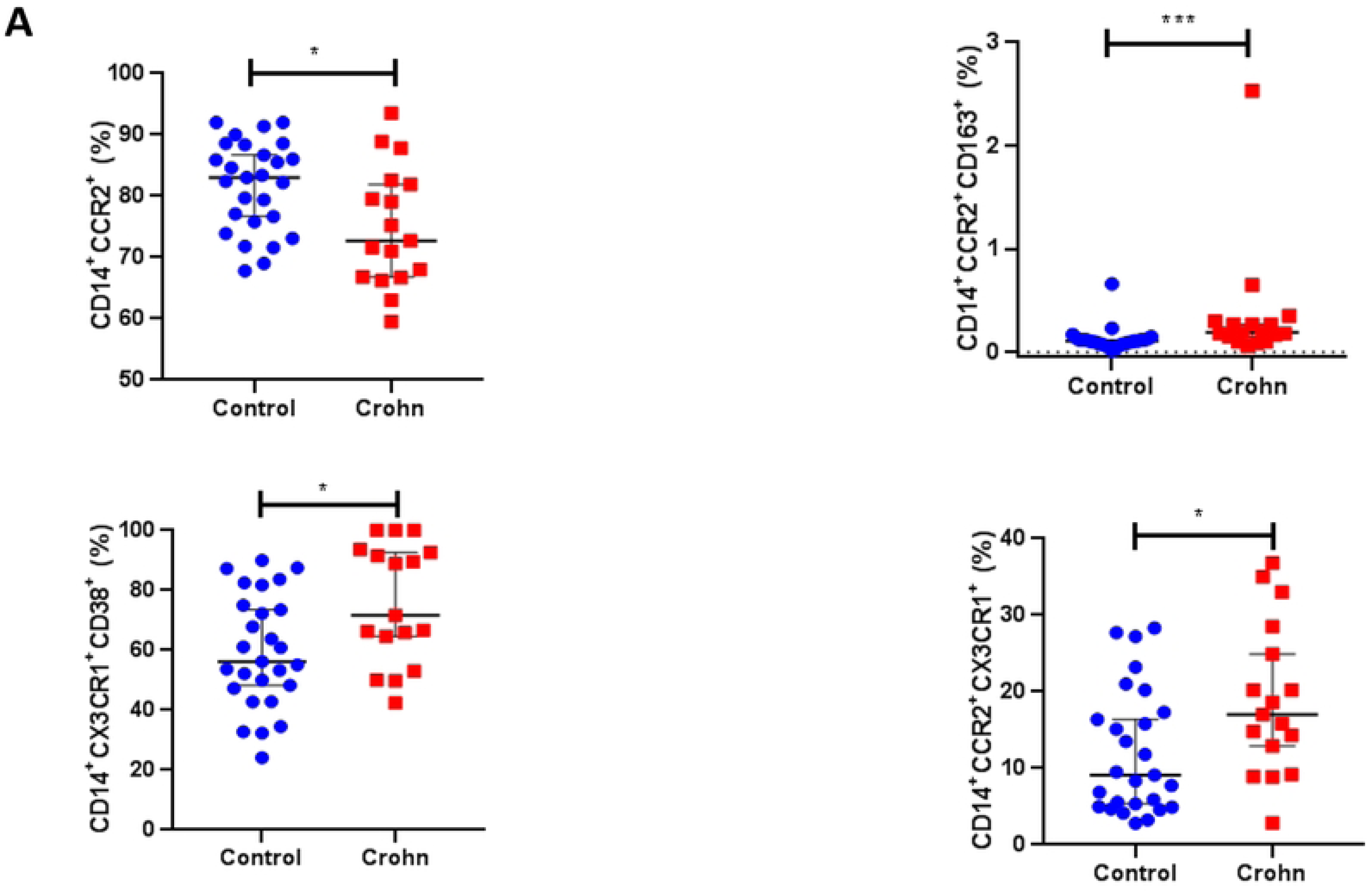
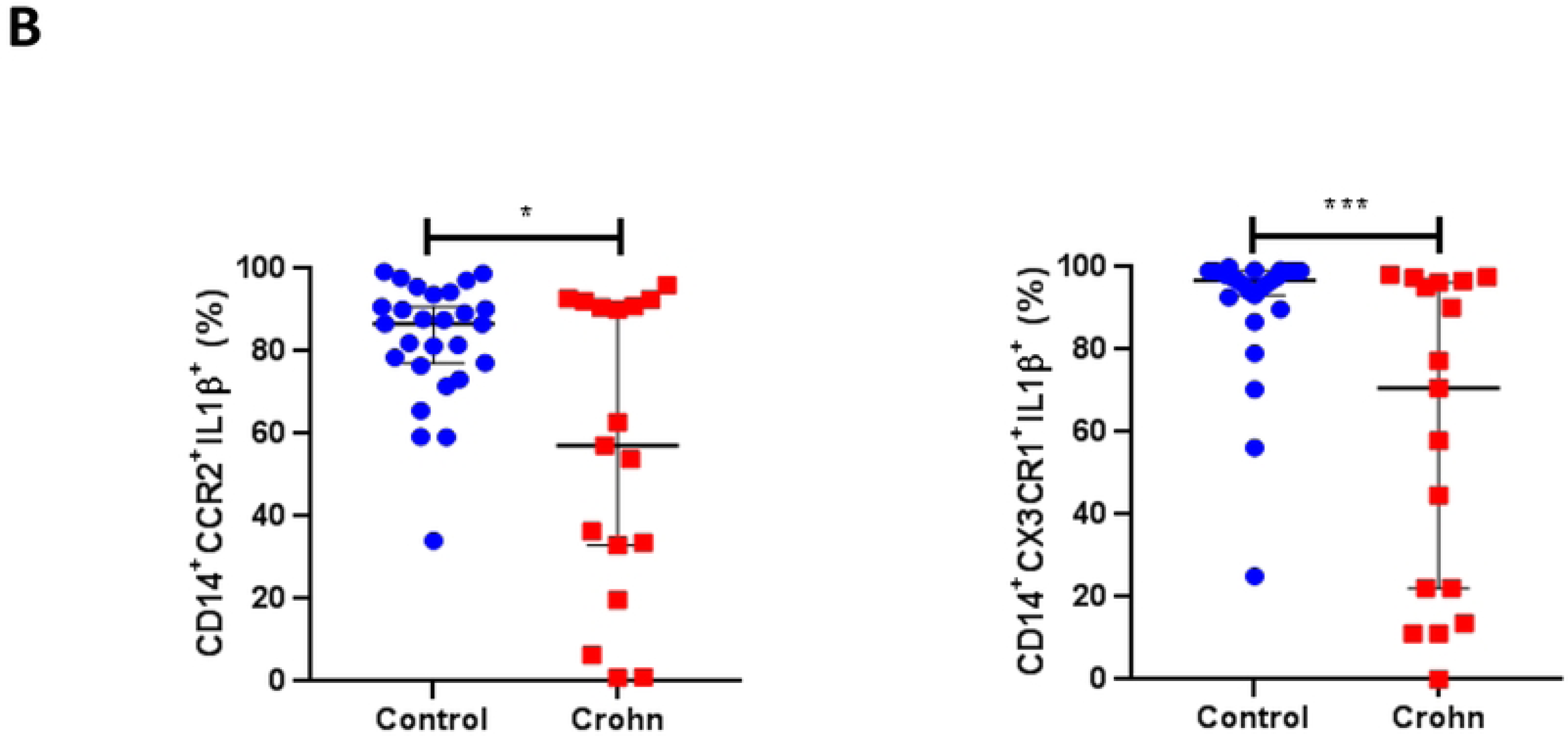
PBMC stimulated with LPS gating on classical monocytes. **(A)** From upper left shows a decrease of CCR2 in Crohn’s patients, an increase in expression of CCR2^+^CD163^+^, CX3CR1^+^CD38^+^ (exhaustion), and CCR2^+^CX3CR1^+^. **(B)** After LPS stimulus, classical monocytes downregulated a secretion of IL-1β and CCR2^+^CX3CR1^+^ in Crohn’s patients.

### After LPS stimulus, CD patients downregulated a secretion of IL-1β

Monocytes are known to be a primary source of IL-1β, even following a single stimulus of TLR2-4 ^[37,38]^. Using LPS stimulus, we observed decreased IL-1β production in classical monocytes expressing CCR2 and CX3CR1 (Fig 3B). These results demonstrate that Crohn’s patients have a decline in crucial proinflammatory cytokine, which could be impaired in a disease relapse.

### Positive correlation on gene expression of TNF-α and NLRP3

We performed a qPCR in total PBMC to verify the gene expression of TNF-α, IL-1β, IL-10, and NLRP3 in both groups. This cytokine and NLRP3 have a role in IBD. The increase of NLPR3, for example, increases the expression of several effectors downstream of this sensor. There were no significant statistical results between healthy subjects and Crohn’s patients. However, we identified a positive correlation between the gene expression of NLPR3 and TNF and IL-10 in healthy subjects (Fig 4A). The exact correlation was observed on Crohn’s patients but just with TNF (Fig 4B). Since the qPCR was done in PBMC, we cannot correlate the above expression with specific myeloid cells such as monocytes. Specifics of all correlations between qPCR and classical monocytes can be found in the supplementary Figure 3A and B.

**Figure 4.**
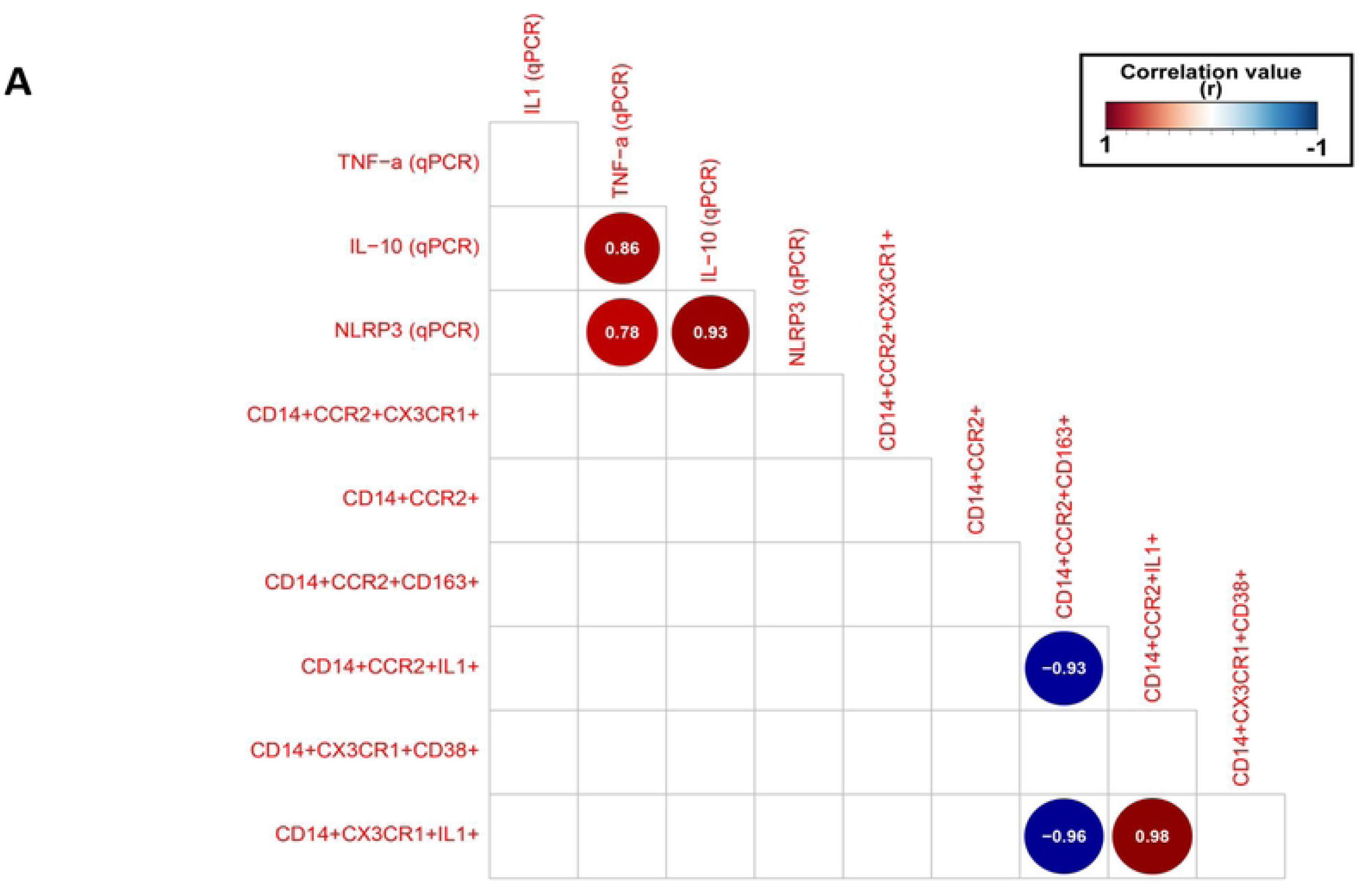
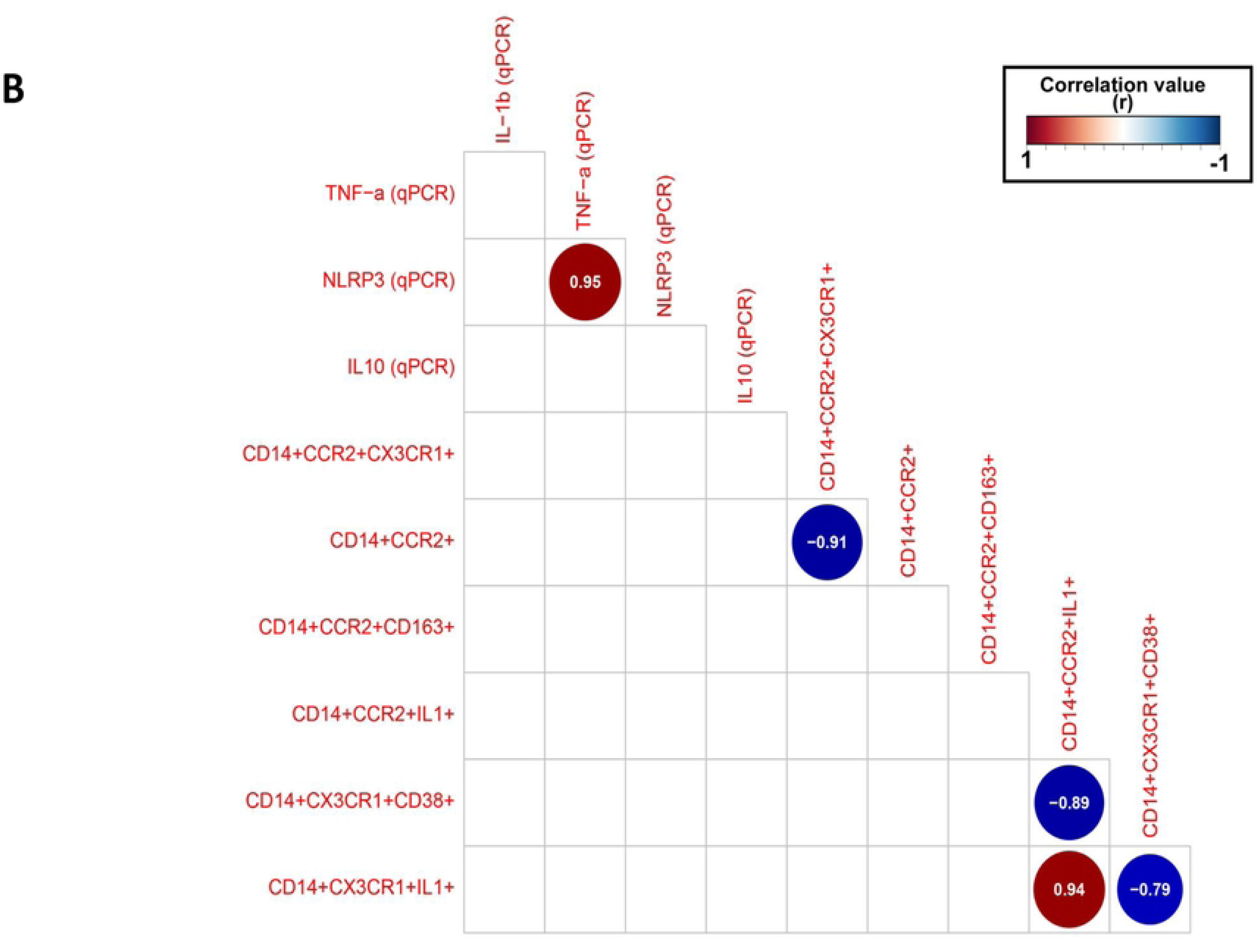
Gene expression correlated to monocytes subsets. **(A and B)** Gene expression correlated to monocyte subsets using Pearson correlation (adjusted p-value ≤0.05). Positive correlation is shown as a red circle, and negative correlation is shown as a blue circle.

### Crohn’s patients show exhaustion homing profile

Classical monocytes expressing CX3CR1 are responsible for migration. Our results show a negative correlation between classical monocytes expressing CX3CR1 secreting IL-1β and the subsets CX3CR1 and CD38 in Crohn’s patients (Fig 4B). That shows the inflammatory state decreases the secretion of IL-1β and promotes classical monocytes to express that a higher concentration of CD38 could promote an exhaustion state.

## Discussion

The incidence of Crohn’s disease is increasing annually. The treatments available around the world provide an improvement in remission and sometimes a disease clearance for many years ^[39]^. Therefore, understanding the innate immune response could give some clues to improve treatments for patients who relapse after discontinuing biologics or do not respond well to the available treatment.

Our results show a decrease in all subsets of monocytes in Crohn’s patients, including an expression of CD163 and CCR2. An overview of a cohort of IBD patients over 19 years shows they were able to demonstrate high monocyte counts, which shows these patients may be more likely to experience relapses ^[40]^. The non-classical monocytes, on the other hand, show an increase in chemokine receptors in Crohn’s patients. Cells that express one or both CCR2 and CX3CR1 are found in inflamed gut tissue of patients with inflammatory bowel disease (IBD), specifically CD^[5]^, thus highlighting chemoreceptors’ importance for inflammatory responses.

The decrease of classical monocytes can stimulate the bone marrow to produce more myeloid cells to compensate for the loss of these cells. Our results show an increase of CD33^+^CD14^−^, showing a possible compensation for the immature cells in the periphery.

The primary sources of IL-1β are monocytes and macrophages following a single stimulus of TLR-2 or TLR-4 ^[41,42]^. To demonstrate the capacity of monocytes function, we stimulated PBMC with LPS. Our cohort of Crohn’s patients shows a decrease in classical monocytes expressing CCR2, CCR2^+^IL-1β^+^, and CX3CR1^+^IL-1β^+^. Chapuy shows the major contributor of IL-1β secretion was CD163 subsets of macrophages in inflamed mucosa of CD ^[20]^. Gareth-Rhys et al. show intestinal myeloid cells in colitis have monocytes, the principal cells of the IL-1-producing population ^[34]^. Even though the results are the opposite of ours, they look at another tissue in other subsets. However, the study from Mitsialis shows an increase of monocytes secreting IL-1β but was in CD active when compared with non-IBD; however, there was no significant difference in CD inactive ^[43]^.

We demonstrated that monocytes are decreased in the circulating blood of Crohn’s patients. After the LPS stimulus, increased CD38 expression and reduced cytokine production were observed compared to healthy individuals, suggesting that CD monocytes display characteristics of exhaustion in such disease.

Cell exhaustion is also reported on monocytes but to a lesser extent. Pradhan, finding repetitive challenger of high dose LPS in murine monocytes derived from bone marrow for 5 days exhibited a pathogenic inflammation of septic monocytes. They identified CD38 as a novel marker for exhausted monocytes ^[24]^. One year later, Naler did a persistent low-dose LPS challenge that led murine monocytes derived from bone marrow to exhaustion, resembling cells observed in septic patients ^[44]^. CD38 expression seems important for cell exhaustion.

Nevertheless, limited data has been published regarding the exhaustion of mononuclear cells in chronic diseases, especially CD. Our results show a characteristic of exhausted monocytes in CD patients; this finding may help us understand the monocyte’s role in CD pathogenesis. We demonstrated that CD14^+^CCR2^+^CX3CR1^+^ and CD14^+^CCR2^+^CD163^+^ cells were increased in the CD patients; this may reflect a skewed profile towards the intestinal tissue in an LPS-induced manner. Along with the presence of CD163, the expression of CCR2 suggests the mobilization of these cells to the intestine^[15–17]^. Furthermore, the expression of CX3CR1 and CCR2 is associated with gut immune cells. Bernardo and colleagues demonstrated an increase in intestinal macrophages expressing CX3CR1^+^CCR2^+^ in patients with IBD compared to healthy individuals. In addition, those macrophages were arising from infiltrating proinflammatory CCR2^+^ monocytes ^[5]^. These findings highlight the presence and importance of CCR2^+^CX3CR1^+^ cells in driving intestinal inflammation and suggest the observed CX3CR1^+^CCR2^+^ and CCR2^+^ populations as being recruited to the inflammation site and associated with antigens in the tissue ^[45]^, thus further contributing to the inflammatory profile.

Although we did not observe differences in gene expression comparison between patients and healthy individuals, the correlation analysis demonstrated in patients a positive correlation between NLRP3 and TNF-α, suggesting an association between inflammatory and inflammasome signaling. The inflammasome pathway has already been correlated with CD. A mutation in the CARD8 gene impedes its downregulation of NLRP3, thus favoring the inflammasome activation, and may also take part as a treatment resistance inducer in patients undergoing monoclonal antibody therapy ^[46]^. Furthermore, Gettler and colleagues performed a genomic analysis demonstrating CD-related gene overexpression in leukocytes. Those genes were predominantly overexpressed in monocytes, thus associating such cell populations with the pathogenesis of this disease ^[47]^, demonstrating that healthy individuals have some differences regarding cytokine gene expression compared to CD patients. The negative correlation between exhausted monocytes and classical monocytes secreting IL-1β sustained the pathogenic inflammation present in CD patients.

A drug that can maintain the monocytes with functional capacity in the bloodstream of Crohn’s patients can improve the lifetime of the patients and diminish their relapsing. It is crucial to note that future studies are required to understand better the source of NAD^+^ and mitochondria function in IBD patients.

## Conclusion

Despite Crohn’s disease being described almost a century ago, it still has some mechanisms and etiology gaps. Our results demonstrated that monocytes subsets may be differentially involved in the pathophysiology. LPS-stimulated PBMC of Crohn’s patients is susceptible to increased cellular exhaustion marker expression and decreased cytokine production, favoring a chronic profile with no disease resolution. Future functional and mechanistic studies in CD are needed to fully elucidate how monocytes correlate with the intestinal tissue and the microbiota.

## Acknowledgments

We thank clinical laboratory staff from Hospital Israelita Albert Einstein for the continuous laboratory support. We warmly thank the volunteers, patients, and their families who contributed to this work.

